# Activating Immune Recognition in Pancreatic Ductal Adenocarcinoma Using Autophagy Inhibition, MEK blockade and CD40 Agonism

**DOI:** 10.1101/2020.11.05.370569

**Authors:** Honglin Jiang, Tristan Courau, Leonard Lupin-Jimenez, Joseph Borison, Alexa J. Ritchie, Aaron T. Mayer, Matthew F. Krummel, Eric A. Collisson

## Abstract

Pancreatic ductal adenocarcinoma (PDA) patients have not yet benefitted from the revolution in cancer immunotherapy due in large part to the dominantly immunosuppressive tumor microenvironment (TME). MEK inhibition combined with autophagy inhibition leads to transient tumor responses in some PDA patients. We find that co-inhibition of MEK (using cobimetinib, COBI) and autophagy (using mefloquine, MFQ), but not either treatment alone, activates the Type I Interferon/STING pathway in tumor cells which in turn reprogram tumor associated macrophages (TAMs) in paracrine to foster an immunogenic switch. This effect is augmented by a CD40 agonist (aCD40). Triple therapy (COBI+MFQ+aCD40) achieved cytotoxic T cell activation in an immunologically “cold” mouse PDA model, leading to enhanced anti-tumor immunity. Collectively, MEK and autophagy co-inhibition coupled with CD40 agonism invokes immuno-reprograming and is an attractive therapeutic approach for PDA immunotherapy development.

## Introduction

Pancreatic ductal adenocarcinoma (PDA) is a leading cause of cancer death in the United States (Rahib et al., 2014). Combination chemotherapy has improved survival in PDA but targeted small molecules, and immunotherapies that have revolutionized the care of many other cancers, have had no meaningful impact in PDA and long-term survival remains low (Ryan et al., 2014). *KRAS* mutations drive >90% of human PDAs (Almoguera et al., 1988). Mutant Ras proteins signal largely down the RAF/MEK/MAPK pathway in PDA (Collisson et al., 2011), but inhibition of this pathway with potent inhibitors of MEK1/2 is compensated for by increases in autophagocytic flux. Inhibiting lysosomal acidification with hydroxychloroquine theraputically synergizes with MEK or ERK inhibition (Bryant et al., 2019; Kinsey et al., 2019) and is currently under clinical investigation (NCT03825289). The combination of MEK inhibition with autophagy inhibition is thought to operate by a tumor cell-autonomous mechanism, without any specific requirement for immunocompetency *per se*.

Reactivation of the immune system to fight pancreatic cancer has been largely unsuccessful to date. Blockade of PD-1/PD-L1, either alone or with CTLA-4 (O’Reilly et al., 2019) or with chemotherapy (Wainberg et al., 2020) were not effective outside of the extremely rare microsatellite-unstable setting (Marabelle et al., 2020) indicating potent immune evasion by PDA cells, a dominant acting immunosuppressive microenvironment (Li et al., 2018), or both.

We sought to explore the functional effects of combined MEK and autophagy inhibition on the PDA immune microenvironment through the study of multiple immune competent, preclinical model systems. We found that co-inhibition of MEK and autophagy led to transcriptional activation of inflammatory cytokines in the cancer cell. These signals affect macrophages polarization to favor a M1-like, antigen-presenting phenotype. This effect was further augmented by CD40 agonism, and lead to T-Cell dependent tumor killing *in vivo*, prolonging overall survival in immunocompetent mouse models of the disease.

## Results

### Mefloquine (MFQ) compares favorably to Hydroxychloroquine (HCQ) to inhibit autophagy in vivo, and synergizes with MEK inhibition in PDA

Inhibition of autophagy in the clinic is currently limited by the onset of action and potency of available agents. Hydroxychloroquine (HCQ) is widely used clinically for the treatment of psoriasis but was developed as an antimalarial medication. Other antimalarial medications, namely mefloquine (MFQ) have superior pharmacokinetic properties to HCQ and may more potently inhibit autophagy in cancer cells (Sharma et al., 2012). To measure autophagy under treatment with HCQ or MFQ both with and without MEK inhibition, we used multiple assessments of autophagic flux. We first confirmed that MEK inhibition led to expected increased autophagosome assembly, in agreement with previous reports (Kinsey et al., 2019), as evidenced by accumulation of LC3B in MiaPaca2 PDA cells (**Figure 1A-B**). Additional treatment with HCQ or MFQ, which inhibit acidification of the lysosome, both effectively led to a further elevated accumulation of LC3B (**Figure 1A-B**). We observed synergistic anti-proliferative effects at MFQ concentrations of 1μM when combined with cobimetinib at 100nM in MiaPaca2 cells (**Figure 1C**). Potent and durable autophagy inhibition is important to prevent MEK inhibitor escape in PDA, so we next tested the duration of autophagy inhibition *in vivo*. We implanted FC1245 mouse PDA cells stably expressing a tandem fluorescence LC3 reporter (mCherry-GFP-LC3) (Kimura et al., 2007) orthotopically and treated tumor-bearing mice with either MFQ or HCQ (**Figure 1D**). Inhibition of lysosome by HCQ or MFQ led to the accumulation non-acidic autolysosomes dually flouresent for red and green LC3. A single dose of MFQ resulted in more GFP positive tumor cells (indicating more potent deacidification of the lysosome) at three and as long as five days after treatment (**Figure 1E-F**). Similarly, treatment of Mia-PaCa2 and PANC-1 reporter cells with either HCQ or MFQ led to the expected increase of GFP positive cells (**Supplementary Fig S1A**). Given these results, we anchored our *in vivo* autophagy inhibition approaches on the MFQ backbone for the remainder of the study.

**Figure 1.**
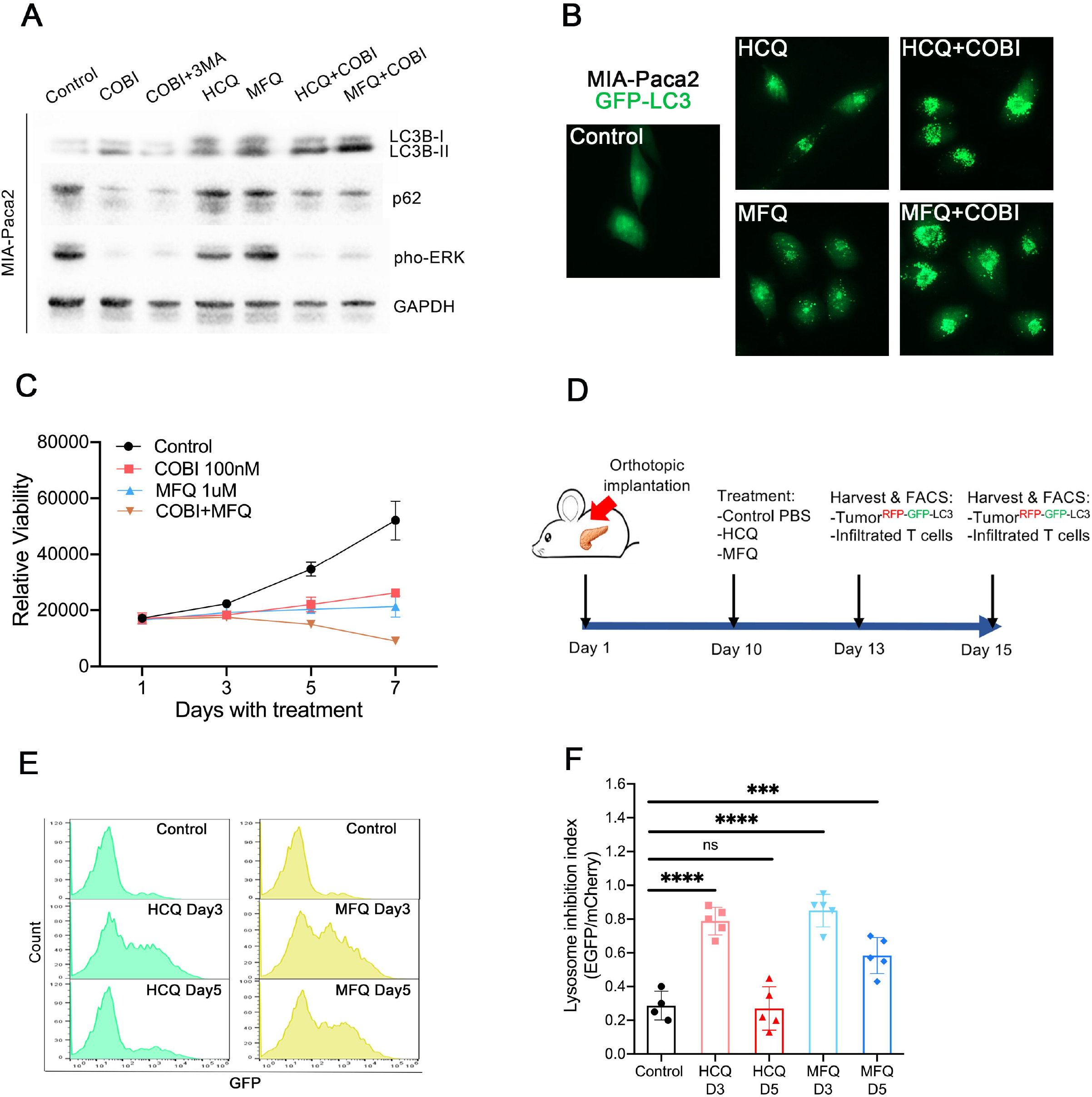
(A) Western blot for LC3B, p62, pho-ERK and total ERK in MiaPaca-2 cells with indicated treatment. (B) Representative images of LC3 expression in MiaPaca-2 cells carrying GFP-LC3 with indicated treatment. (C) Viability of MiaPaca-2 cells with indicated treatment. Error bars indicate ±SD. (D) Scheme of implantation, dose and harvest schedule for E and F. (E) Representative histogram assessing GFP expression of FC1245 tumors stably expressing mCherry-GFP-LC3. (F) Quantification of lysosome inhibition measured by the percentage of EGFP/mCherry. Error bars represent mean ± SEM, n = 3mice/group and analyzed by two sample t test. ***p < 0.001.

### Combined MEK and autophagy inhibition activates STING-Type I-IFN pathway in PDA cells

The cytokine secretion profile of PDA cells is driven by Kras and plas an important role in immune evasion (Hou et al., 2020; Muthalagu et al., 2020; Wang et al., 2019). To evaluate the effects of MEK and autophagy inhibition on cytokine secretion by PDA cells, we examined transcription of a panel of inflammatory cytokines in syngeneic PDA orthotopic tumors (FC1245) from mice treated with MFQ, Cobimetinib, or both. We harvested the tumors after treatment, sorted out tumor cells and found that the Type I interferons *Ifn-α/β* were dramatically upregulated under combined MEK and autophagy inhibition. Type II IFNs (IFN-γ) and other cytokines examined were not induced by co-treatment (**Figure 2A**). *In vitro* treatment of previously described immunologically “cold” PDA cell lines (6694C2)(Li et al., 2018) confirmed that either MFQ or HCQ combined with inhibition of MEK increased Type 1 IFN transcription, lending generalizability to this treatment approach in PDA (**Figure 2B**). We next used an IFN-α/β sensitive reporter cell line B16-blue (Rees and Lowy, 2018) to quantify the amount of functional IFN protein present in the media of mouse PDA cells treated with the indicated drugs. We found that combined MEK and autophagy inhibition produced upregulation of functional Type I IFN proteins (**Figure 2C**). The innate immune STING-IFN pathway plays a critical role in antiviral defense and cancer. We found that treatment with cobimetinib and MFQ led to STING pathway activation as evidenced by increased expression of cGAS and pho-IRF3 (**Figure 2D**). A similar STING-Type I-IFN pathway activation was observed with *in vitro* treatment in a panel of human PDA cells, as evidenced by increased mRNA transcription of IFNs, STATs and IRFs (**Figure 2E**). We next treated 6694C2 PDA cells with different concentrations of cobimetinib or MFQ, either alone or in combination, and assessed drug synergy/antagonism by the Loewe Additivity method (**Figure 2F**)(Chou, 2010). We observed synergistic anti-proliferative effects was of MFQ and MEK combinations at MFQ concentrations in the 5–15μM range when combined with cobimetinib in the range of 1–10nM (**Figure 2F**). Interestingly, much lower levels were needed to increase production of Type I IFNs indicating that lower, more clinically tolerable doses might be sufficient to elicit this *in vivo* effect (**Figure 2G**). We conclude from these experiments that combined MEK and autophagy inhibition cooperate to activate STING-Type I-IFN pathway in PDA cancer cells at doses below those required for their cytostatic/cidal effects.

**Figure 2.**
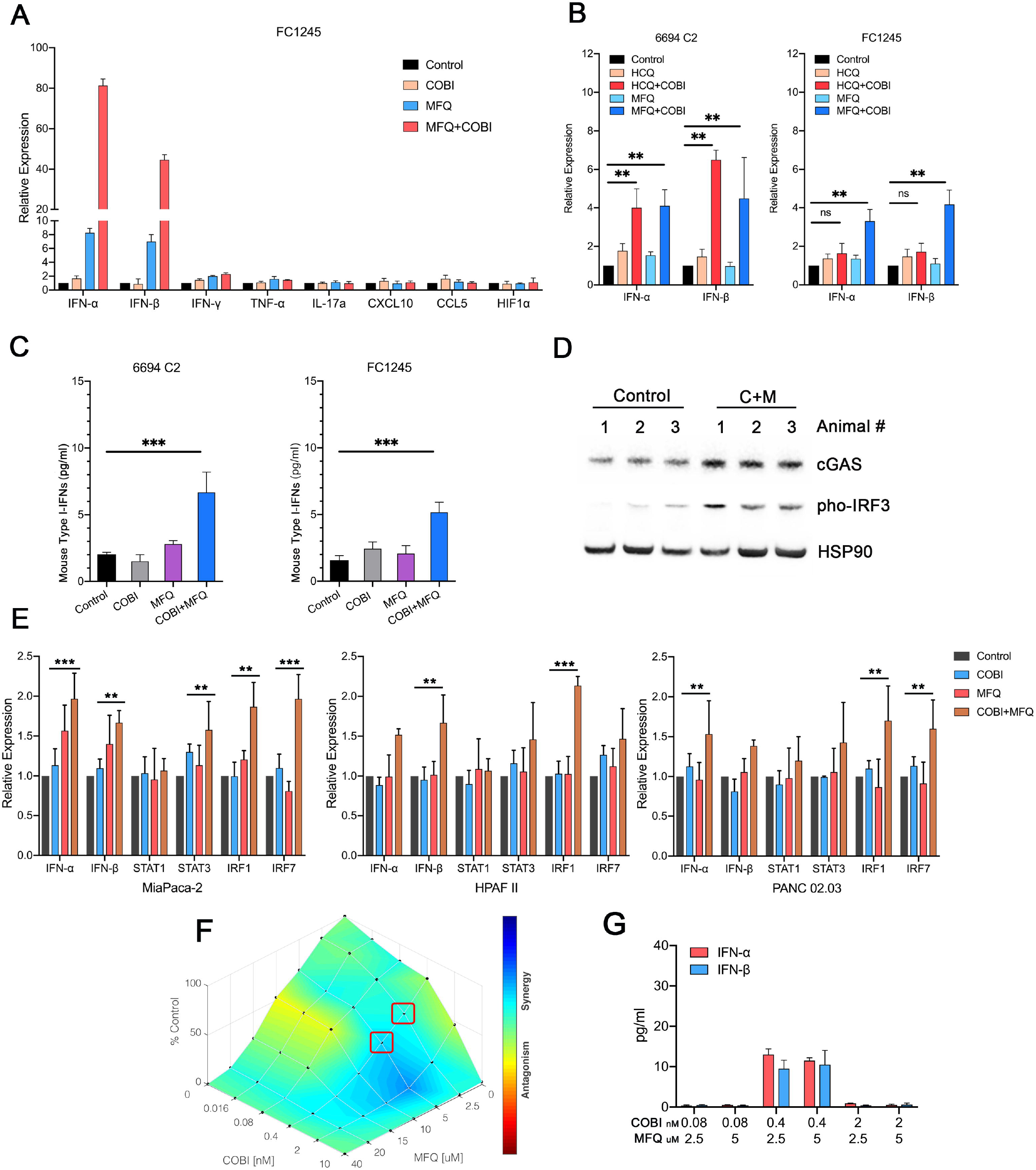
(A) Quantitative PCR analysis of a panel of cytokines in FC1245 tumor cells freshly isolated from FC1245 orthotopic engraftments with indicated treatment. Error bars represent mean ± SEM. (B) Quantitative PCR analysis of IFN α/β in 6694C2 and FC1245 cells receving indicated treatment *in vitro*. Error bars represent mean ± SEM. Data are analyzed by two sample t test. **p < 0.01. (C) The amount of IFN proteins present in the media of mouse PDA cells (6694C2 and FC1245 cells) treated with the indicated drugs quantified by IFN-α/β sensitive B16-blue reporter cell line. Error bars represent mean ± SEM. Data are analyzed by two sample t test. ***p < 0.001. (D) Western blot for cGAS and pho-IRF3 expression in whole tumor lysates after C+M treatment and vehicle. (E) Quantitative PCR analysis of IFN α/β, STAT1/3 and IRF1/7 in MiaPaca-2, HPAFII and PANC02.03 cells with indicated treatment. Error bars represent mean ± SEM. Data are analyzed by two sample t test. **p < 0.01, ***p < 0.001. (F) 6694C2 cells were treated for 48h as indicated with cobimetinib and MFQ and analyzed for cell viability. Synergy graphs were generated utilizing Combenefit Software. (G) 6694C2 cells treated with the combination of cobimetinib and MFQ and IFN-α/β production measure by ELISA assay. Error bars represent mean ± SEM.

### CD40 agonist synergizes with COBI/MFQ to inhibit PDA progression

Type I IFNs directly or indirectly affect multiple immune cell types, including monocytes, antigen-presenting cells, NK cells and lymphocytes (Gessani et al., 2014) (Hervas-Stubbs et al., 2011; Schiavoni et al., 2013). Certain tumor-associated macrophages (TAMs) and certain populations of dendritic cells both serve as antigen presenting cells (APCs) and promote cytotoxic T cells recruitment in the tumor microenvironment. These APCs become fully “licensed” after CD40 ligand engagement (Beatty et al., 2011). As such, we hypothesized that the doublet of COBI/MFQ might functionally synergize with CD40 activation to improve the control of tumor outgrowth in an IFN-1 and T cell dependent manner.

To explore the therapeutic potential of adding activating CD40 ligand to the doublet regimen, we first orthotopically implanted C57BL/6J mice with syngeneic, immunologically “cold” 6694C2 cells (Li et al., 2018). We began treatemetn five days after implantation using low doses of COBI plus MFQ followed by aCD40 mAb in an intermittent dosing schedule as shown in **Figure 3A.** Tumor growth was significantly decreased in COBI/MFQ/aCD40 treated mice compared to either COBI/MFQ or aCD40 mAb mono-treated mice (**Figure 3B-C, Supplementary Figure S1B**). In addition, treatment of established 6694C2 tumors on day 10 with COBI/MFQ/aCD40 achieved significant regression and enhanced survival, including apparent cures (**Figure 3D-E**). We subsequently rechallenged mice that were cured of the primary tumor with the triplet of COBI/MFQ/aCD40 with a second implantation of 6694C2 tumor cells (this time subcutaneously) 40 days later and observed a failure of engraftment despite robust growth of the same cells implanted in treatment-naive control Black6 animals (**Figure 3F**, **Supplementary Figure S1C**). Similar results were seen with a second syngeneic PDA cell line (FC1245) also derived from a C57BL/6J KPC mouse (**Figure 3G-H**).

**Figure 3.**
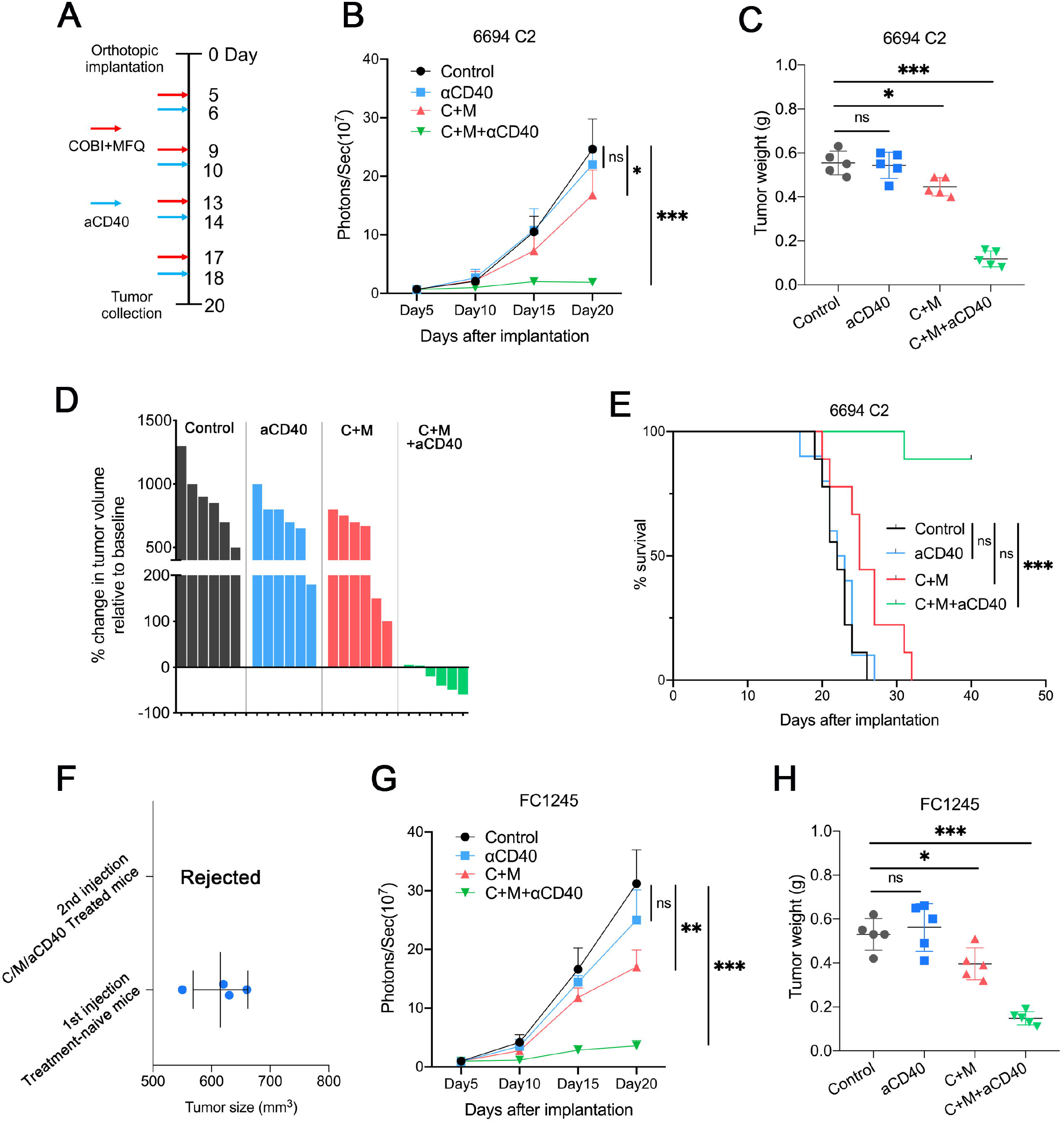
(A) Schematic illustration of the dose schedule after C57BL/6J mice were orthotopically engrafted with mouse PDA cells. (B) Mean tumor growth and (C) tumor weights for mice bearing 6694C2-fLuc tumors with indicated treatments. n=5 mice/group. Data are analyzed by one-way ANOVA (B) and two sample t test (C). *p < 0.05, ***p < 0.001. (D) Change in 6694C2 tumor volume on day 20 compared to the start of treatment on day 7, representative of 6 mice per group. (E) Kaplan-Meier survival curves of mice engrafted with 6694C2 tumors with indicated treatment. Minimum n = 8 mice per group, log-rank test, ***p < 0.001. (F) Tumor sizes for mice after second subcutaneous injection of 6694C2 cells >40 days after primary orthotopic implantation. Treatment-naïve WT C57BL/6J mice receiving first time subcutaneous injection served as positive controls. (G) Mean tumor growth and (H) tumor weights for mice bearing FC1245-fLuc tumors with indicated treatments. n=5 mice/group. Data are analyzed by one-way ANOVA (G) and two sample t test (H). *p < 0.05, **p < 0.01, ***p < 0.001.

### COBI/MFQ/aCD40 combination therapy triggers immune activation in the PDA tumor microenvironment

To profile the immune microenvironment (TME) in an unbiased manner, we next performed mass cytometry (CyTOF) of 6694C2 tumours after treatment with COBI/MFQ/aCD40, aCD40 alone, or vehicle to investigate the distribution of immune cells in TME upon the treatments (**Figure 4A**). Globally, COBI/MFQ/aCD40 induced striking changes in clusters of lymphocytes, macrophages and monocytes (**Figure 4B-D, Supplementary Figure S2A**). CD3^+^ T cells comprised ~5% of all CD45^+^ cells in vehicle-treated mice, increased slightly under treatment with aCD40 monotherapy and were dramatically elevated by the addition of COBI/MFQ. CD4 regulatory T cells were significantly decreased after COBI/MFQ/aCD40 therapy (**Figure 4E**). The combination of COBI/MFQ/aCD40 slightly enhanced the infiltration of monocytes but had no effects on the overall prevalence of TAMs and conventional dendritic cells (cDCs)(**Figure 4F**). In agreement with earlier reports that the STING-IFNs pathway can prime strong effector activity in NK cells (Marcus et al., 2018), we also observed that the expression of CD16/32 and CD11b was significantly increased on these NK cells (**Figure 4G**). Histologic analysis of the tumors confirmed a remarkable increased infiltration of both CD8^+^ T cells and CD4^+^ T cells in the tumor (**Figure 4H**). Furthermore, we observed expected decrased collagen deposition and tumor cell proliferation after the treatment (**Supplementary Figure S2B**).

**Figure 4.**
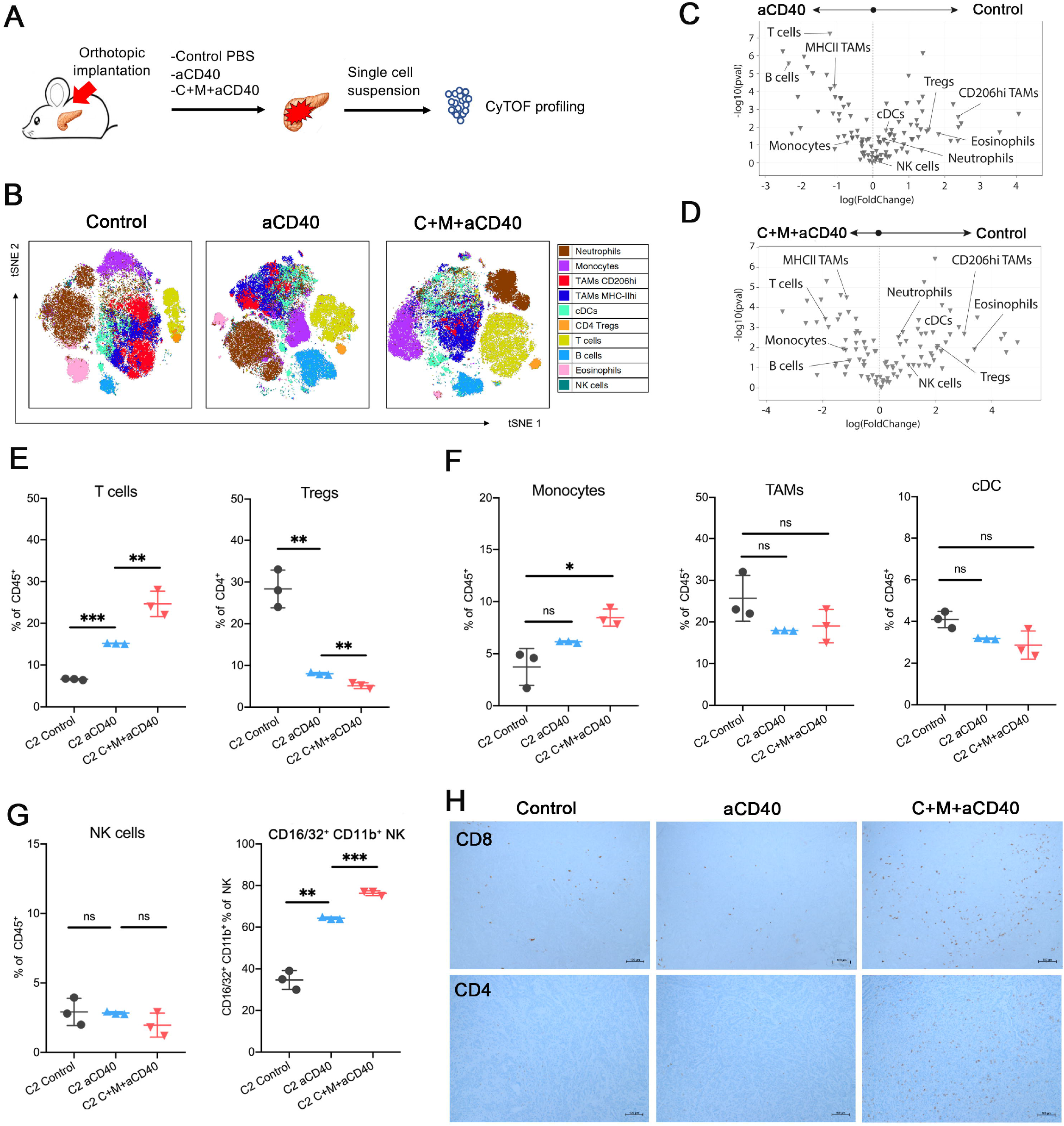
(A) Schematic illustration of mouse PDA infiltrating immune cells analyzed by CyTOF. (B) t-SNE projection showing the distribution of immune populations in orthotopic tumors with indicated treatment. Each population is identified by a distinct color. (C-D) Volcano plots depicting differential immune population percentages for aCD40 or COBI/MFQ/aCD40 treatment versus control. (E) Quantification of total T cells among CD45+ and Treg cells among CD4+ T cells. (F) Quantification of monocytes, total tumor associated macrophages and and conventional dendritic cells among CD45+ cells. (G) Quantification of total natural killer cells among CD45+ and CD16/32+ among NK cells. (H) IHC of orthotopic engratments stained with CD8 and CD4. E,F and G: n=3 mice/group. Data are analyzed by two sample t test. *p < 0.05, **p < 0.01, ***p < 0.001.

CD8^+^ T cells, increased with therapy, and were essential for the triple therapy’s antitumor effects, as evidenced by anti-CD8+ neutralizing antibody treatments administered prior to the initial treatment with COBI/MFQ/aCD40 (**Supplementary Figure S4B**). Given the increased infiltration of CD8^+^ T cells upon treatment, we investigated whether checkpoint inhibitor could enhance tumor growth inhibition. However, mice treated with COBI/MFQ/aCD40/anti-PD-L1 showed similar response rates as mice with COBI/MFQ/aCD40 (**Supplementary Figure S4C-D**). Taken together, our data reveals that a durable anti-tumor immunological program is activated upon the triple therapy in mice bearing “cold” PDA tumors.

### Triplet of COBI/MFQ/aCD40 Promotes Immunogenic Macrophage Polarization

Although CyTOF data showed similar prevalence of tumor associated macrophages (TAMs) and conventional dendritic cells (cDCs) with COBI/MFQ/aCD40 compared to COBI/MFQ or aCD40 alone (**Figure 4F**), macrophage expression of MHC-II, considered as part of the M1 phenotype, was increased in aCD40 monotherapy and the addition of COBI/MFQ increased its expression. By contrast, COBI/MFQ/aCD40 concomitantly downregulated the expression of CD206, considered as part of the M2 phenotype (**Figure 4C-D, Figure 5A**). Moreover, consistent with our CyTOF results, COBI/MFQ/aCD40 combination resulted in upregulated transcription of genes related to M1 polarization (*Cxcl10, Tnfa, Il15, Nos2*) of macrophages, whereas M2-associated transcription factors were found downregulated (*Il10, Mrc1, Ym1*; **Supplementary Figure S3A**).

**Figure 5.**
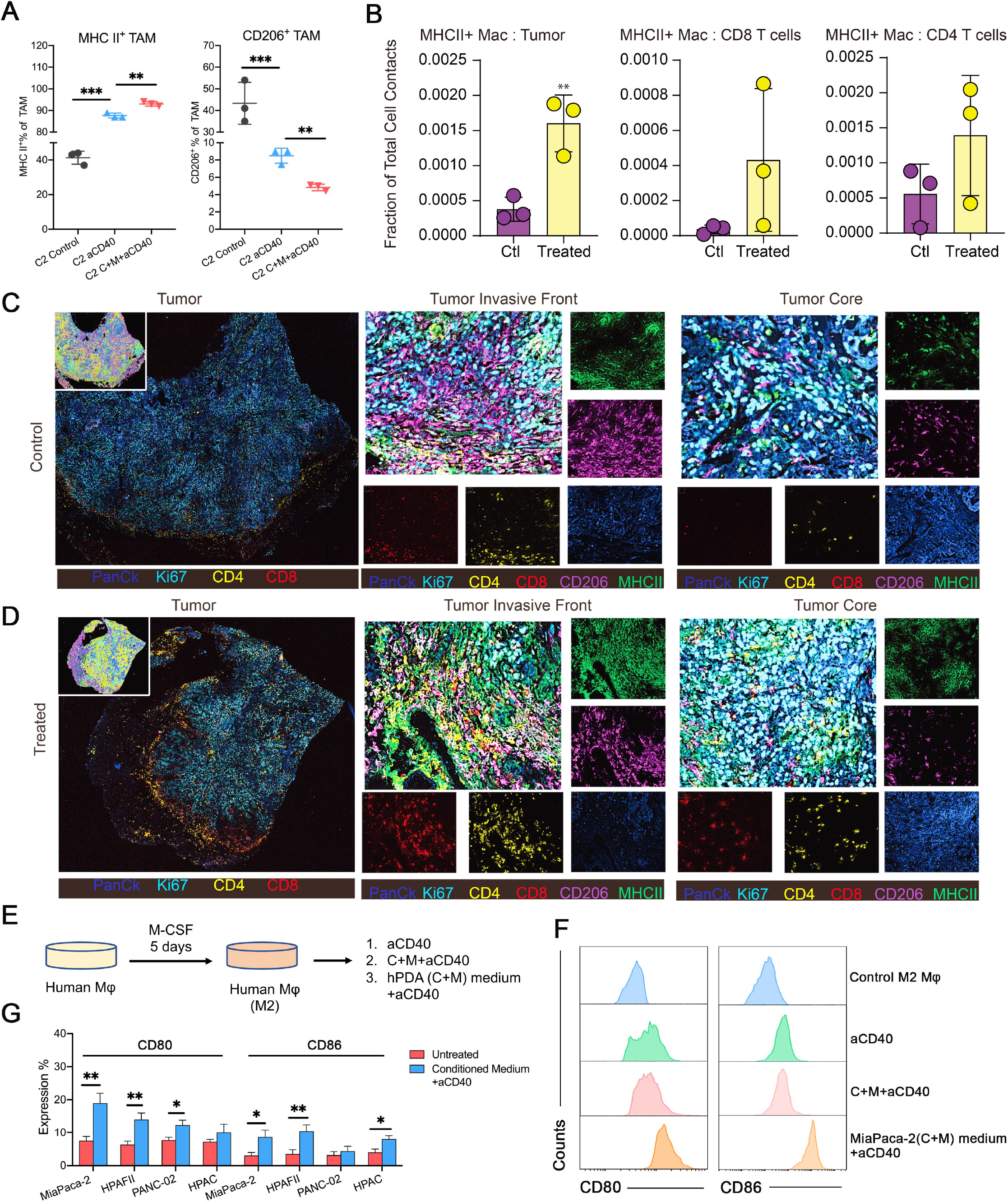
(A) Quantification of MHCII+ M1-like and CD206+ M2-like macrophage subsets among all TAMs. (B) Quantification of cell-to-cell interaction scores between MHCII+ TAMs and tumor cells or CD8+ T cells or CD4+ T cells. Error bars represent mean ± SEM. (C-D) Representative multiplexed immunofluorescence images of COBI/MFQ/aCD40 treated and non-treated tumors determined by CODEX shows immune exclusion and immune infiltration. Zoom panel shows individual immune compartments, blue=anti-PanCK, cyan=anti-Ki67, yellow=anti-CD4, red=anti-CD8, magenta =anti-CD206, green=anti-MHCII. (E) Schematic illustration of conditioned media transfer experiments in vitro. (F) Representative histogram assessing staining for CD80 (left) and CD86 (right) by flow cytometry in human macrophages treated directly with indicated drugs and conditioned media from MiaPaca-2 cells. (G) Quantification of CD80 and CD86 expression in human macrophages treated with vehicle and conditioned media from MiaPaca-2, HPAFII, PANC02 and HPAC cells. Error bars represent mean ± SEM, analyzed by two sample t test. **p < 0.01, ***p < 0.001.

We next performed CODEX multiplexed inmmunofluorescence imaging (Goltsev et al., 2018) in order to better characterize the intratumoral architecture at a single-cell and cellular interaction level. MHCII and CD206 expression tightly colocalized in macrophages, and the T cells often were excluded along these boundaries in untreated tumors. In contrast, most of MHCII expression did not colocalize with CD206 in the treated tumors. T cells were also well infiltrated throughout many areas of the tumor (**Figure 5C-D**). Furthermore, we observed increased pairwise cell-cell interactions between CD4^+^ or CD8^+^ T cells and MHC II^+^ macrophages in COBI/MFQ/aCD40 treated tumors compared to untreated tumors, suggesting the treatment triggered enhanced antigen presentation with functionally important intercellular / microenvironmental sequelae (**Figure 5B**).

To demonstrate the communication between adenocarcinoma cells and macrophages, we next performed conditioned media transfer experiments *in vitro*. We exposed primary isolated human macrophages to 5 days of M-CSF to induce the M2 state, and then exposed these M2-polarized macrophages to media from four human PDA cell lines (MiaPaca2, HPAFII, Panc-02, and HPAC) which had been pretreated with COBI and MFQ for 72 hours. We also added aCD40 to the conditioned media on human macrophages (**Figure 5E**). The conditioned media + aCD40 in all cases increased macrophage activation (as evidenced by CD80 and CD86 upregulation) (**Figure 5F-G, Supplementary Figure S3B**). In contrast, direct treatment of macrophages with cobimetinib, MFQ and aCD40 had moderate effect on macrophage repolarization *in vitro* (**Figure 5F**). Strikingly, depletion of macrophages *in vivo* in the 6694C2 model via injection of anti-F4/80 antibodies abrogated the response to COBI/MFQ/aCD40 treatment, indicating that the anti-tumor immunity in these settings is, in addition to CD8 T cells, also dependent on macrophages (**Supplementary Figure S4B**). Together, these data suggest the combined regimens of COBI/MFQ/aCD40 reprogrammed TAMs toward an immunogenic M1-like phenotype through a paracrine mechanism involving PDA cells’ production of diffusible interferons.

### CD40 agonist enhances COBI/MFQ efficacy in a STING-Type I-IFN dependent manner

To ascertain the requirement of Type I IFNs in mediating COBI/MFQ/aCD40 tumor inhibition, we treated mice with Type 1 IFN receptor blocking antibody in addition to COBI/MFQ/aCD40 and observed failure of tumor control by the triple therapy (**Figure 6A-B**). We also confirmed reduction of M1 macrophages by flow cytometry after this blockade (**Figure 6C**). Next, we aimed at determining the extent to which the synergistic antitumor effect of combined COBI/MFQ/aCD40 therapy is mediated through tumor cell cGAS-mediated STING activation. To accomplish this, we depleted either cGAS or STING in 6694C2 cells using shRNA and estimated knockdown efficiency by Western blot analysis (**Figure 6D**). Knockdown of cGAS or STING, dramatically decreased tumor growth inhibition relative to sh-control expressing tumors treated with the triplet regimen (**Figure 6E, Supplementary Figure S4A**), confirming the vital role of the cGAS/STING pathway in COBI/MFQ/aCD40-mediated antitumor response in PDA model. We harvested non-fully rejected tumors from this experiment to perform flow cytometry and observed that depletion of either STING or cGAS fully abrogated the macrophage M1 polarization (**Figure 6F**) and the infiltration of CD8^+^ T cells (**Figure 6G**).

**Figure 6.**
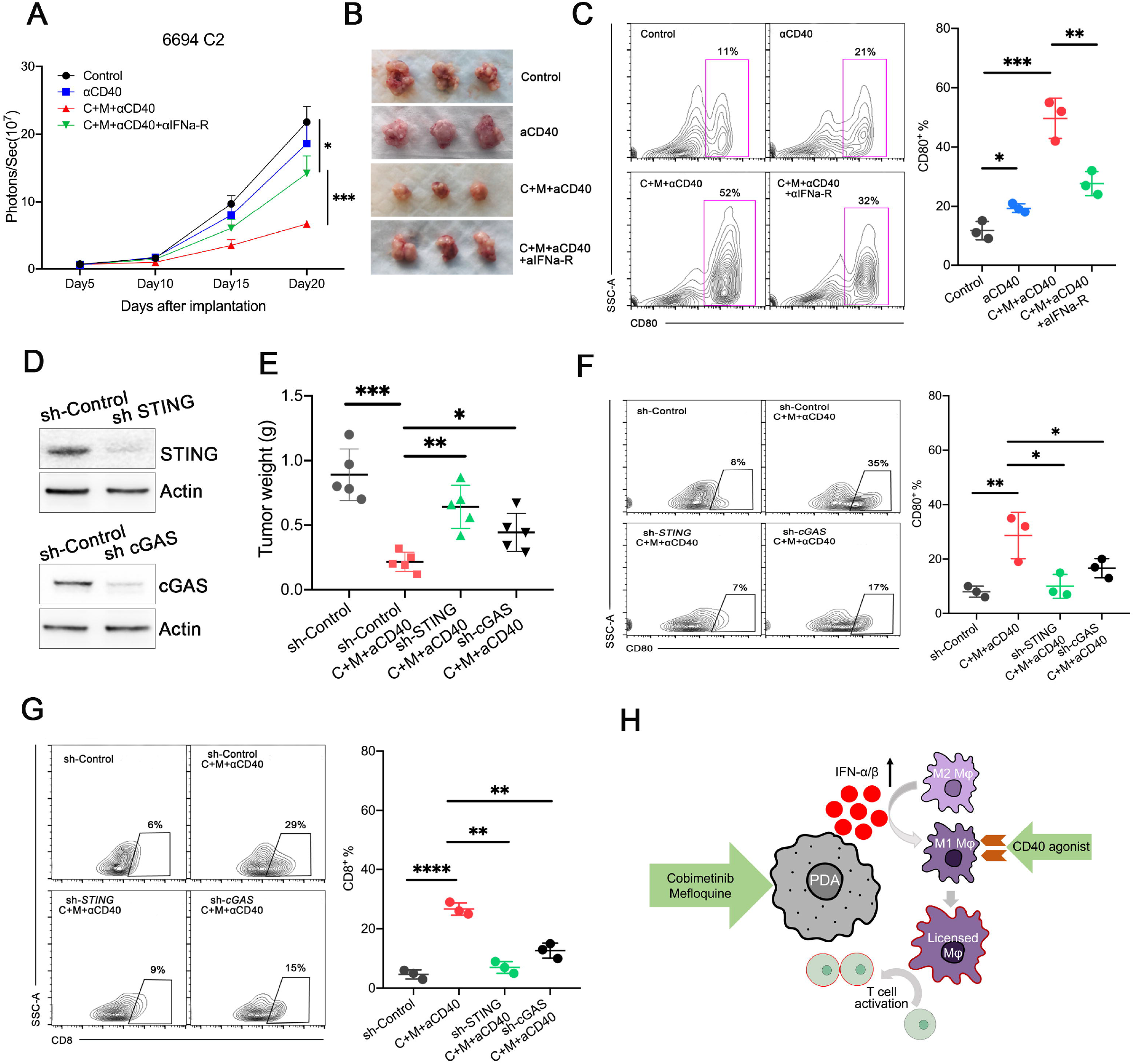
(A) Mean tumor growth and (B) representative pictures of tumors for mice bearing 6694C2-fLuc tumors with indicated treatments. n=5 mice/group. Data are analyzed by two sample t test. *p < 0.05, ***p < 0.001. (C) Quantification of CD80+ M1-like macrophage subsets in 6694C2 tumors with indicated treatment. n=3 mice/group. Data are analyzed by two sample t test. *p < 0.05, **p < 0.01, ***p < 0.001. (D) Western blot for STING and cGAS in 6694C2 cells after indicated shRNA knockdown. (E) Tumor weights for mice bearing 6694C2-shCon tumor, 6694C2-sh-cGAS tumor and 6694C2-sh-STING tumor with COBI/MFQ/aCD40 treatment. Error bars represent mean ± SEM, n=5mice/group and analyzed by two sample t test. *p < 0.05, **p < 0.01, ***p < 0.001. (F) Quantification of CD80+ M1-like macrophage subsets in 6694C2-shCon tumor, 6694C2-sh-cGAS tumor and 6694C2-sh-STING tumor with COBI/MFQ/aCD40 treatment. Error bars represent mean ± SEM, n=3 mice/group and analyzed by two sample t test. *p < 0.05, **p < 0.01. (G) Quantification of CD8+ T cells in 6694C2-shCon tumor, 6694C2-sh-cGAS tumor and 6694C2-sh-STING tumor with COBI/MFQ/aCD40 treatment. Error bars represent mean ± SEM, n=3 mice/group and analyzed by two sample t test. **p < 0.01, ***p < 0.001. (H) Working model. Combined MEK and autophagy inhibition activated the Type-1 Interferon/STING pathway in PDA tumor cells which in turn reprogramed tumor associated macrophages (TAMs) towards an immunogenic phenotype. This effect was augmented with therapeutic CD40 agonism to convert an immunologically exclusive tumor into one with abundant T cell infiltration.

## Discussion

PDA treatment has not yet improved from the addition of immune checkpoint inhibitors, largely due to: 1) a paucity of neoantigens arising from a relatively low tumor mutational burden, and 2) an immunoinhibitory microenvironment which dominantly antagonizes potentially tumoricidal T cells. In this study, we aimed to improve anti-cancer immunity in PDA. Using a series of mutant KRAS-driven mouse model of PDA, we found that treatment with a CD40 agonist combined with autophagy and MEK inhibition altered multiple cellular subpopulations in the tumor immune microenvironment, bringing it from an immuno-suppressive towards a T cell-active milieu, supporting immunological control of tumor outgrowth. Combined blockade of autophagy and MEK activated the STING-Type I-IFN pathway in PDA cells resulting in improved antigen presentation and subsequent robust infiltration of cytotoxic T cells mediated by M1-like TAMs (**Figure 6H**).

Previous studies have highlighted autophagy’s complex role in facilitating oncogenic RAS-driven proliferation. Lock et al. demonstrated that the autophagy pathway is required for efficient secretion of multiple cytokines including IL-6, which promotes tumor cell invasion (Lock et al., 2014). Kinsey et al. and Bryant et al. also unveiled a protective role of autophagy in response to inhibition of RAS-RAF-ERK signalling to preserve tumor cell fitness in PDA (Kinsey et al., 2018),(Bryant et al., 2019). Yamamoto et al. then identified an autophagocytic processes as a driver of immunoevasion in the tumor cell by the posttranslational downregulation of MHC-I presentation and T cell surveillance (Yamamoto et al., 2020).

Successful therapeutic interventions maximize efficacy and minimize toxicity. In the present study, we show that sub-cytotoxic doses of combined autophagy and MEK inhibitors triggered activation of the STING-Type I IFN pathway in PDA cells. In addition to low efficacy, regimens in previous studies adding immune therapeutic agents to established chemotherapy backbones seemed poorly tolerated, limiting the ability to improve upon them. Encouragingly, the intermittent administration of COBI/MFQ (two doses per week) plus CD40 agonism was sufficient for successful *in vivo* antitumor effects. Although Byrne et al. (Byrne and Vonderheide, 2016) argue that innate immune sensing pathways are not required for CD40 stimulation to trigger adaptive T cell activation, we find that the primed STING-Type I IFN signaling greatly boosted CD40-agonist-mediated anti-tumor immunity by a series of loss-of-function *in vivo* studies.

Pharmacologic CD40 agonism is actively being investigated for an improved antitumor response (Beatty et al., 2011; Byrne and Vonderheide, 2016; Lin et al., 2020; Long et al., 2016). Although single agent aCD40 mAb was found to be able to modulate myeloid subpopulations in TME, it failed to achieve significant anti-tumor immunity. We therefore examined the therapeutic benefits of aCD40 mAb as an partner with COBI/MFQ therapy in PDA models. Compared to aCD40 mAb monotherapy, we found that the triplet of COBI/MFQ/aCD40 therapy facilitated a much greater degree of TAMs shifting toward an M1 phenotype as well as T cell infiltration. Furthermore, spatial distribution analysis by multiplexed immunofluorescence directly demonstrated increased cell-to-cell interaction frequencies between CD4 or CD8 T cells and MHC II-expressing TAMs, strongly supporting the hypothesis that enhanced antigen presenting is required in COBI/MFQ/aCD40 mediated anti-tumor immnunity. Morever, there appears to be additional activation of NK cells due to the triplet treatment which is in agreement with previous findings (Muthalagu et al., 2020) and warrant further investigation.

In summary, we show MEK inhibition with concurrent autophagy blockade to lead to a STING-dependent increase in Type I IFN production from mouse and human KRAS-driven PDA cells. The activation of STING-Type I IFN signaling can be augmented with therapeutic CD40 agonism to convert an immunologically exclusive tumor into one with abundant T cell infiltration. Our findings suggest that CD40 agonism may clinically synergize with MEK and autophagy inhibition to augment anti-tumor immunity in PDA patients.

## Supplementary figure legends

**Figure S1**

(A) Autophagic flux was assessed by flow cytometry in Mia-PaCa2 and PANC02 cells expressing mCherry-GFP-LC3AFR cells following 48h of indicated treatment. Error bars represent mean ± SEM, data are analyzed by two sample t test, *p < 0.05, **p < 0.01, ***p < 0.001.

(B) Representative pictures of tumors for mice bearing 6694C2-fLuc tumors with indicated treatments.

(C) Representative pictures of mice after second subcutaneous injection of 6694C2 cells >40 days after primary orthotopic implantation (Top). WT C57BL/6J mice receiving first time subcutaneous injection served as positive controls (Bottom).

**Figure S2**

(A) Representative CyTOF dot plots showing gating strategy defining pan T cells, MHCII+ vs CD206+ TAMs, CD16/32+/CD11b+ NK cells and Treg cells.

(B) Images of HE, trichome staining and IHC staining with Ki67 in 6694C2 tumors with indicated treatments.

**Figure S3**

(A) Quantitative PCR analysis of CXCL-10, TNFα, IL-15, NOS2, IL-10, MRC1, YM1 and Arginase1 in macrophages freshly isolated from 6694C2 tumors with indicated treatments. Error bars represent mean ± SEM. Data are analyzed by two sample t test, *p < 0.05, **p < 0.01, ***p < 0.001.

(B) Representative histogram assessing staining for CD80 and CD86 by flow cytometry in human macrophages treated with vehicle oronditioned media from HPAFII, PANC02 and HPAC cells.

**Figure S4**

(A) Representative pictures of tumors for mice bearing 6694C2-shCon tumor, 6694C2-sh-cGAS tumor and 6694C2-sh-STING tumor.

(B) Tumor weights and representative pictures of tumors for mice bearing 6694C2-fLuc tumors with indicated treatments. n=5 mice/group. Data are analyzed by two sample t test. *p < 0.05, ***p < 0.001.

(C) Tumor weights and representative pictures of tumors for mice bearing 6694C2-fLuc tumors with indicated treatments. n=5 mice/group. Data are analyzed by two sample t test. ***p < 0.001.

(D) Quantification of T cells in tumors for mice bearing 6694C2-fLuc tumors with indicated treatments. n=3 mice/group. Data are analyzed by two sample t test. ***p < 0.001.

**Figure S5**

(A) Representation of gating strategy used in CyTOF to indentify various populations.

(B) Representation of gating strategy used in CODEX multiplexed imaging analysisto indentify various populations.

## Acknowledgments

This work was supported by NIH, NCI Grants R01 [CA178015, CA222862, CA227807, CA239604, CA230263], U24 [CA210974], U54 [CA224081] grants (EAC). Content does not reflect the views of the Department of Defense, American Cancer Society, National Cancer Institute or National Institutes of Health.

## Authors Contributions

Conception and design: H.J., E.A.C.

Development of methodology: H.J., J.B., A.T.M.,

Data Acquisition: H.J., T.C., J.B., A.T.M., E.A.C.

Analysis and interpretation of data: H.J, T.C., A.T.M. M.F.K., E.A.C.

Writing and review of the manuscript: H.J., E.A.C.

Study supervision: M.F.K., E.A.C.

## Declaration of Interests

E.A. Collisson is consultant at Takeda, Merck, Loxo and Pear Diagnostics, reports receiving commercial research grants from Astra Zeneca, Ferro Therapeutics, Senti Biosciences, Merck KgA and Bayer and stock ownership of Tatara Therapeutics, Clara Health, BloodQ, Guardant Health, Pacific Biosciences and Exact Biosciences

## STAR METHODS

### KEY RESOURCES TABLE

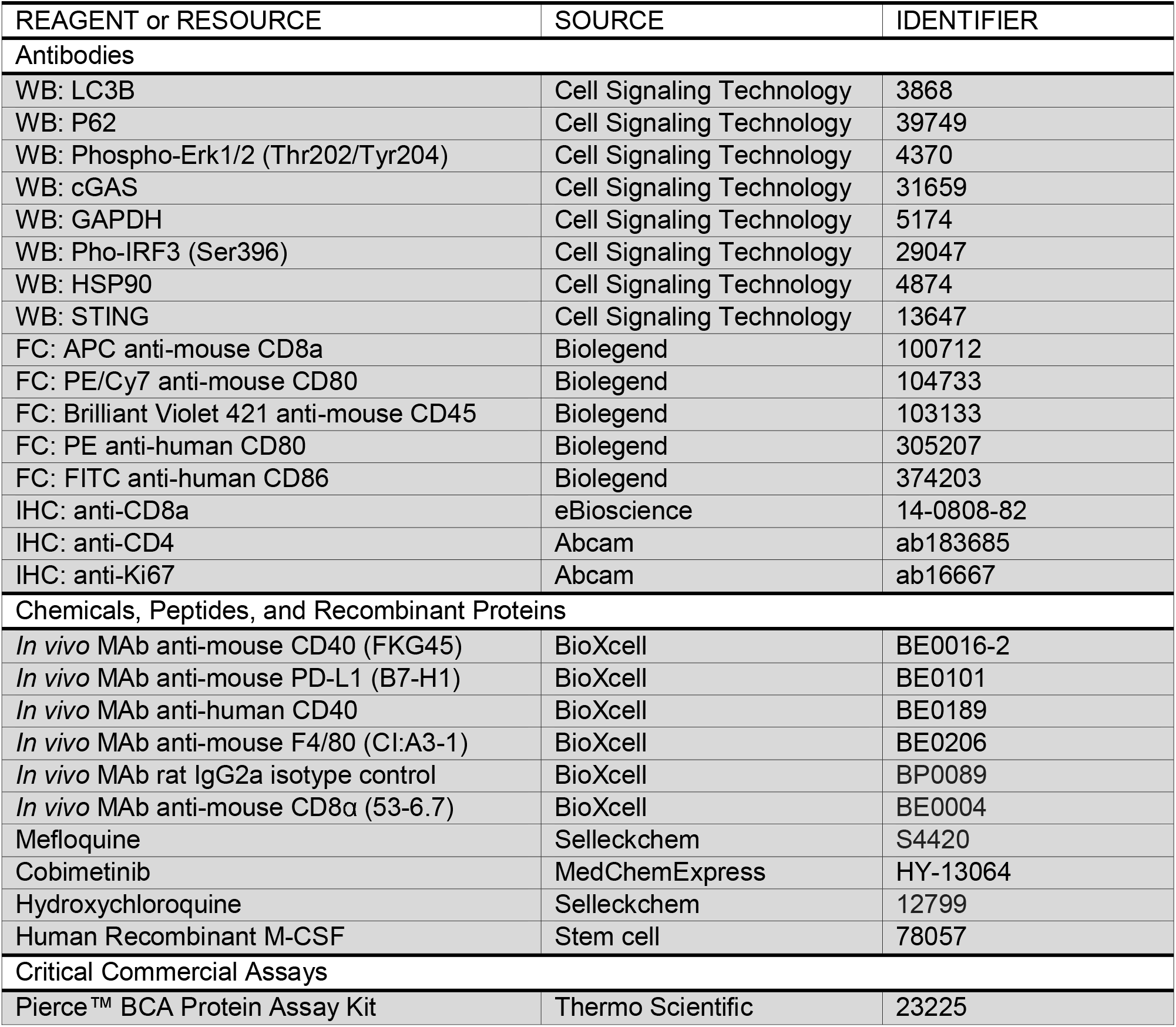
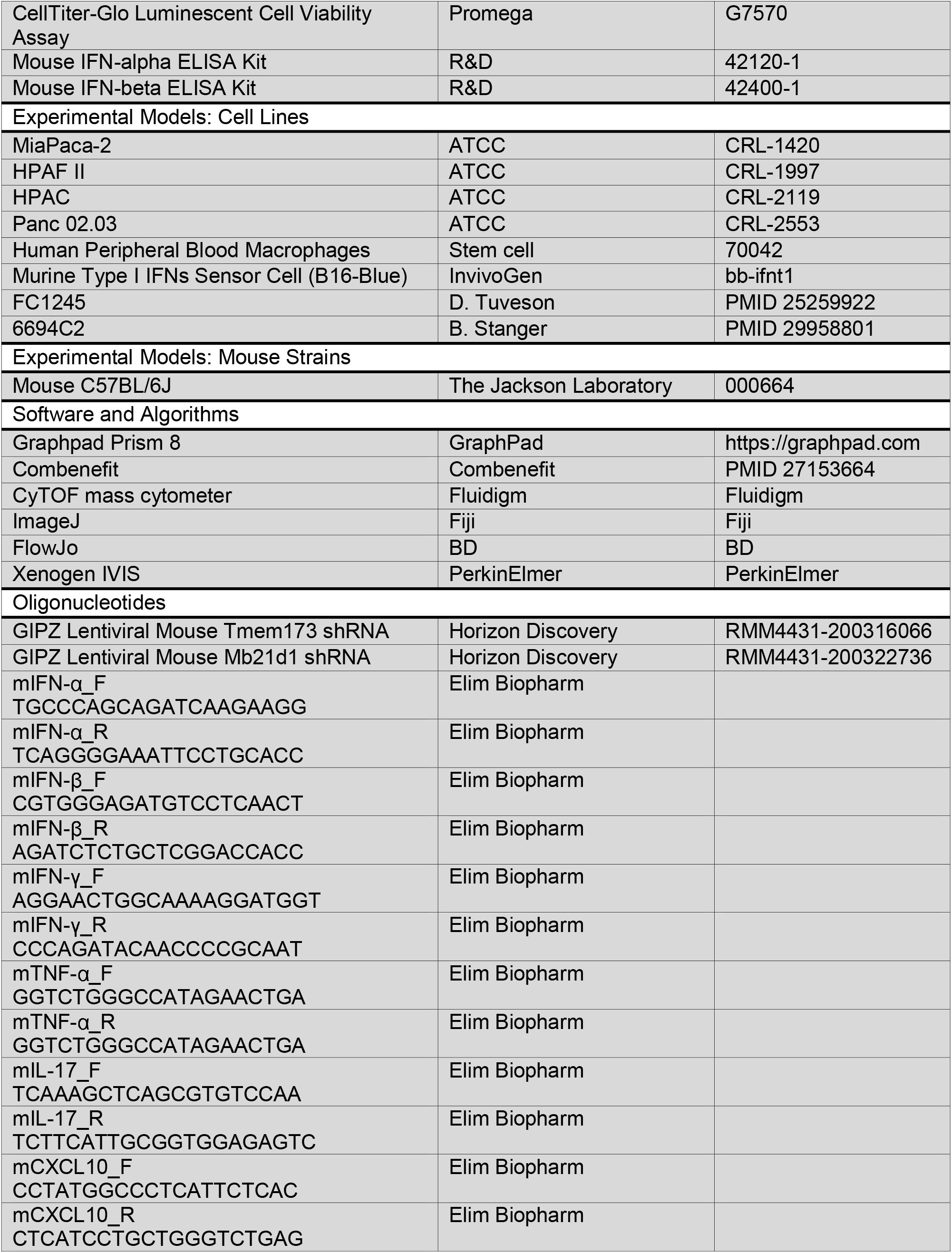
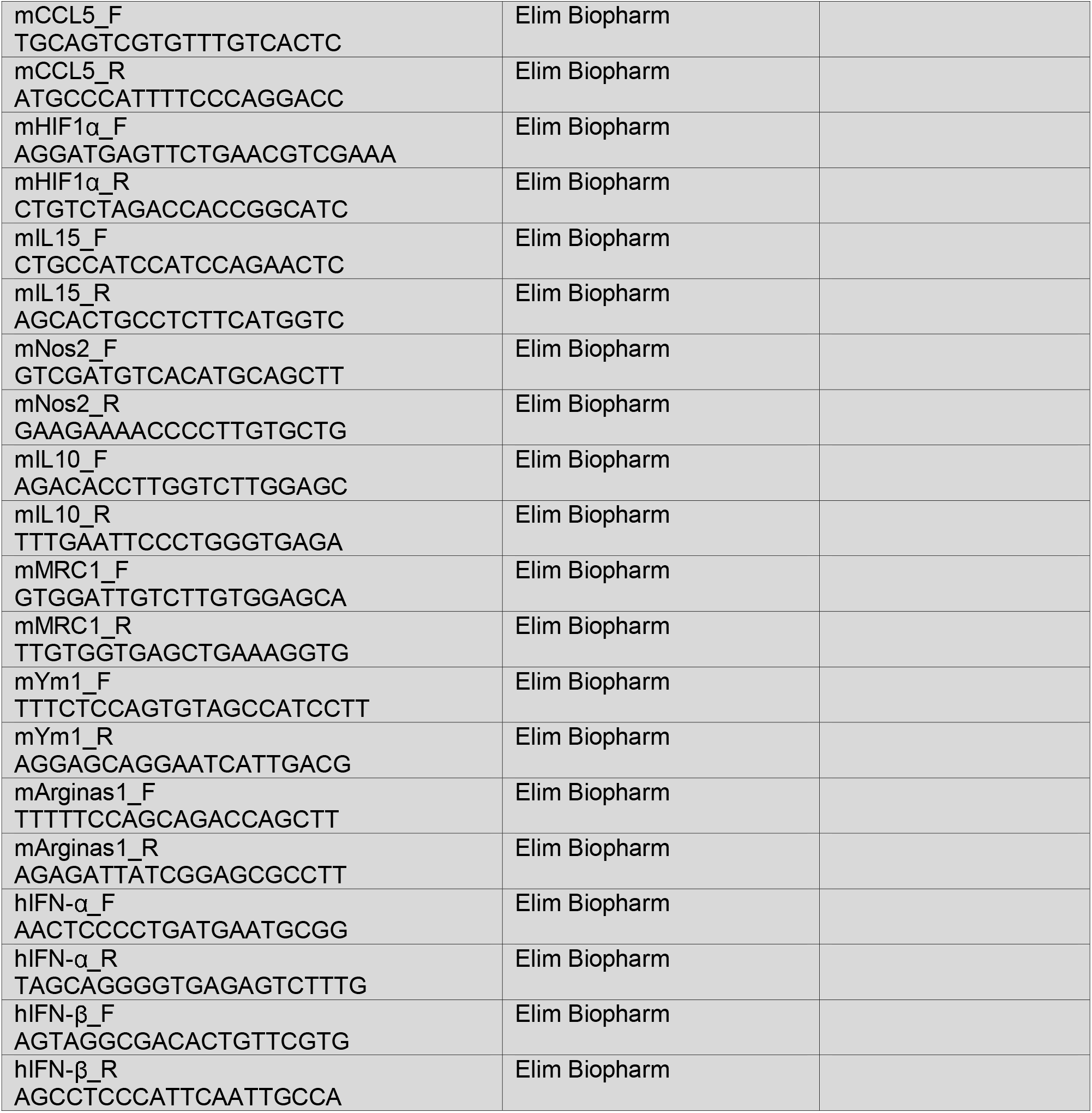

### LEAD CONTACT AND MATERIALS AVAILABILITY

Requests for additional information about the manuscript or for resources and reagents should be directed to and will be fulfilled by the Lead Contact, Eric Collisson. All unique reagents generated in this study are available from the Lead Contact without restriction.

### EXPERIMENTAL MODEL AND SUBJECT DETAILS

#### Cell lines

MiaPaca-2, HPAF II, HPAC and Panc 02.03 are from American Type Culture Collection. 6694C2 cells were kindly provided by Dr. Ben Stanger and FC1245 cells were kindly provided by Dr. David Tuveson. Cells were maintained at 37°C in a humidified incubator at 5% CO2. Cells were grown in appropriate media as recommended by ATCC, and supplemented with 10% fetal bovine serum (Gibco) and 1% penicillin/streptomycin (Gibco). All cell lines tested were negative for mycoplasma contamination.

#### Mice

##### Orthotopic pancreas implantation

6-7 weeks old male C57BL/6J mice were purchased from Jackson lab. All animal experiments were conducted in the AAALAC accredited University of California, San Francisco in accordance with all applicable local requirements, including approval by the IACUC.

## METHOD DETAILS

### Animals and *In Vivo* Procedures

C57BL/6 mice were purchased from Jackson Lab. For orthotopic pancreas implantation, 6-7 weeks old mice were administered intra-pancreatic injections of 6694C2 (provided by Dr. Ben Stanger) or FC1245 (provided by Dr. David Tuveson) PDA cells derived from KPC mice. Cells were transduced *in vitro* with a lentiviral vector encoding a firefly luciferase. Cells were suspended in PBS with 50% Matrigel (BD Biosciences) and 5X10^5^ cells for 6694C2-fLuc and 1000 cells for FC1245-fLuc tumor cells were injected into the body of the pancreas via laparotomy. Bioluminescent imaging (Xenogen IVIS) was performed twice a week to monitor tumor growth. Mice were sacrificed ~3 weeks later and tumors harvested for analyses. Alternatively, mice bearing parental 6694C2 tumors were monitored by ultrasound (Vevo 2100) and analyzed in tumor progression and survival experiments. Treatment was then initiated on day 5 after implantation with vehicle control (PBS/isotype control IgG2a mAb), aCD40 mAb(FKG45) at 100ug followed by cobimetinib at 5 mg/kg (i.p) and MFQ at 50mg/kg (oral) every 4 days. In some experiments, CD8+ T cells or macrophages were depleted with each of clone 53-6.7 and clone CI:A3-1 neutralizing mAbs on day 4 and repeated every 5 days.

### Single Tumor Cell Preparation

Single cell suspensions of PDA tumors were prepared for CyTOF analysis as follows. Briefly, tumors were placed in cold RPMI-1640 medium with Collagenase IV (4 mg/mL) and DNase I (0.1mg/mL), then minced to sub-millimeter pieces. Tissues were then incubated at 37°C for 25 min with gentle shaking every 5 min. Specimens were filtered through a 70 mm mesh and centrifuged at 500 g for 5 min at 4°C. Cells were then resuspend in RPMI-1640 medium with 2% serum for analysis.

### Mass Cytometry (CyTOF)

Mass cytometry was performed as described elsewhere (Allen, Hiam et al. Nat Med 2020). Briefly, conjugations of mass cytometry antibodies with metal isotopes were done using the Maxpar^®^ conjugation kit (Fluidigm) according to manufacturer’s protocols and each antibody was titrated to define its optimal staining concentration. Each sample, initially stained with cisplatinium, fixed with 3.2% PFA and frozen at −80°C on day of harvesting, was thawed and barcoded by mass-tag labelling with distinct combinations of stable Pd isotopes in 0.02% saponin in PBS before further pooling and staining. For staining, cells were first resuspended in cell-staining media (Fluidigm) containing metal-labeled antibodies against CD16/32 for 5 min at room temperature to block Fc receptors, followed by the addition of a cocktail containing surface markers antibodies in a final volume of 500μL for 30 min at room temperature. Cells were then permeabilized with methanol for 10 min at 4 °C, washed and incubated with a cocktail containing intracellular markers antibodies in a final volume of 500μL for 30 min at room temperature. Cells were finally stained with 191/193Ir DNA intercalator (Fluidigm) diluted in PBS with 1.6% PFA 48h prior to data acquisition. For acquisition, cells were washed and resuspended at 1M/mL in deionized water + 10% EQ four element calibration beads (Fluidigm) and run on a CyTOF mass cytometer (Fluidigm). We acquired an average of 1-3 × 10^5^ cells per sample, consistent with generally accepted practices in the field. After data collection, we used the Premessa pipeline (https://github.com/ParkerICI/premessa) to normalize data and deconvolute individual samples. We then manually gated the individual FCS files using FlowJo (BD) according to the gating scheme described in supplementary material. tSNE dimensionality reductions plots were also created using FlowJo software (BD) on total CD45+ events and using all the markers present in the stain panel. To find significant parameters that best explain the variance between groups, we then used a two-sample t-test to calculate the fold change and p-value associated to each manually-gated parameters, plotted them as a classical volcano plot and highlighted on it the populations presented in the tSNE plots. Representation of the gating strategy was shown in **Supplementary Figure S5A**.

### CODEX Multiplexed Imaging Analysis

A 23-plex custom CODEX mouse panel was developed and validated (Enable Medicine) utilizing purified, carrier-free antibodies conjugated to unique DNA oligonucleotides. CODEX staining and imaging was performed on pancreatic mouse tumors utilizing this panel following the manufacturer’s protocol (Akoya Bioscience). Raw fluorescent TIFF image files were processed, deconvolved and background subtracted, and antibody staining was visually assessed for each biomarker and tissue region using the ImageJ software (Fiji, version 2.0.0). TIFF hyper stacks were segmented based on DAPI nuclear stain, pixel intensities were quantified, and spatial fluorescence compensation was performed, which generated comma-separated value (CSV) and flow cytometry standard (FCS) files for downstream analysis. Cell types were enumerated via manual gating (Immune Atlas) based on canonical marker expression values and were confirmed via visual overlay of the cell populations on the immunofluorescent images utilizing the multiplex analysis viewer (MAV, Akoya Biosciences). Voronoi diagrams were created based on the spatial coordinates of the cell types and utilized to mathematically compute adjacent “cell-cell” contacts. Representation of the gating strategy was shown in **Supplementary Figure S5B**.

### Cell Viability Assays

3000-5000 cells optimized for each cell line were seeded on day 1, drugs were added on day 2 (using DMSO normalized to 0.1%), and the cell viability was determined using CellTiter-Glo (Promega) on day 4. Viability curves were generated using GraphPad Prism 6.

### Histological Analysis

HE staining and immunohistochemistry were performed on 4-μm-thick sections of 4% paraformaldehyde-fixed and paraffin-embedded tissues. Tail sections were decalcified by incubation of trimmed paraffin blocks for 10-15 min on paper towels soaked in 1 N HCl. Epidermis thickness was measured by Zeiss AxioImager microscope. Tumors sections were stained with anti-CD8a(14-0808-82), anti-CD4(ab183685) and anti-Ki67(ab16667).

### B16 Cell Mediated IFN Reporter Assay

B16-Blue cells (InvivoGen) were used to measure Type-I IFNs in collected media per manufacturer’s instructions. Briefly, cells were cultured in RPM-1640 medium containing 5% FBS, 1% penicillin/streptomycin and zeocin (100 μg/ml). Cells were then seeded at 50,000 cells per well in a 96 well. Conditioned media from FC1245 or 6694C2 cells with indicated treatment were added to B16 cells for 24 hours alongside IFNα (0–1000 U/mL) to generate a standard curve. QUANTI-Blue was then added in 1:1□v/v ratio for following 24 hs. Media was transferred into a 96 well plate and measured by plate reader.

### Western Blotting

Tumor tissue and organ tissues was flash frozen in liquid nitrogen and homogenized in M-PER lysis buffer (Thermo Scientific) plus Halt™ protease and phosphatase inhibitor cocktail (Thermo Scientific). Protein concentrations were determined with the Pierce BCA Protein Assay Kit (Thermo Scientific), and extracts were loaded onto NuPAGE Bis–Tris SDS gels and immunoblots were visualized by LiCOR Odyssey system.

### Quantitative PCR (qPCR) Analysis for Gene Expression

RNA was prepared from cultured tumor cells or sorted cells from implanted tumors using RNeasy Mini Kit (QIAGEN). cDNA was generated using High-capacity cDNA Reverse Transcription Kit (Bio-Rad). qPCR analysis was performed using TB Green Premix Ex Taq reagent (TaKaRa) and QuantStudio5 qPCR platform, and results were normalized to the expression of 18S rRNA. Primer sequences utilized for qPCR were listed in the key resources table.

### *In Vitro* Synergy Assay

To evaluate synergy *in vitro*, cells were seeded into 96-well plates in complete medium, cultured overnight, and then treated in triplicate with cobimetinib or MFQ, either alone or in various combinations. After 48 hours, cells were assayed using CellTiter-Glo (Promega) according to the manufacturer’s protocol. Luminescence was quantified and analyzed with Combenefit software (Loewe model).

### ELISA-Based Interferon Determination

Supernatants from cells were collected at the indicated times. IFN-α and IFN-β were analyzed by ELISA kits (R&D Systems) with manufacturer’s instructions.

### Lentivirus Transduction For Sh-RNA Mediated Knockdown

For stable and lentivirally transfected shRNA-based knockdown experiments, viruses were generated in HEK 293T cells transfected with lentiviral packaging vectors along with vectors expressing pGIPZ-shRNA using Fugene6 (Promega). Two distinct hairpins were chosen for the experiments. Tmem173(STING) shRNA (RMM4431-200316066) and Mb21d1(cGAS) shRNA (RMM4431-200322736) are purchased from Horizon Discovery. Viral supernatant collected from confluent monoculture was filtered and used to infect 6694C2 cells. A total of 0.5□×□10^6^ cells was seeded in one well of a 6-well chamber and allowed to grow overnight. The following day, cells were incubated with a 1:2 mixture of growth medium and viral supernatant collected from HEK 293T cells. Polybrene was added at 8□μg/ml.

## QUANTIFICATION AND STATISTICAL ANALYSIS

Statistical tests were performed using GraphPad Prism 7.0. Two-sided two-sample t-tests were used for comparisons of the means of data between two groups. One-way ANOVA was used for comparisons among multiple independent groups. Significance of overall survival was determined using Kaplan-Meier survival curve with log-rank analysis. For animal studies, animals were randomized before treatments, and all animals treated were included for the analyses.

## Notes

### Competing Interest Statement

The authors have declared no competing interest.

