## Supplementary figures and images for "Activating Immune Recognition in Pancreatic Ductal Adenocarcinoma Using Autophagy Inhibition, MEK blockade and CD40 Agonism"

### Supplemental Fig1

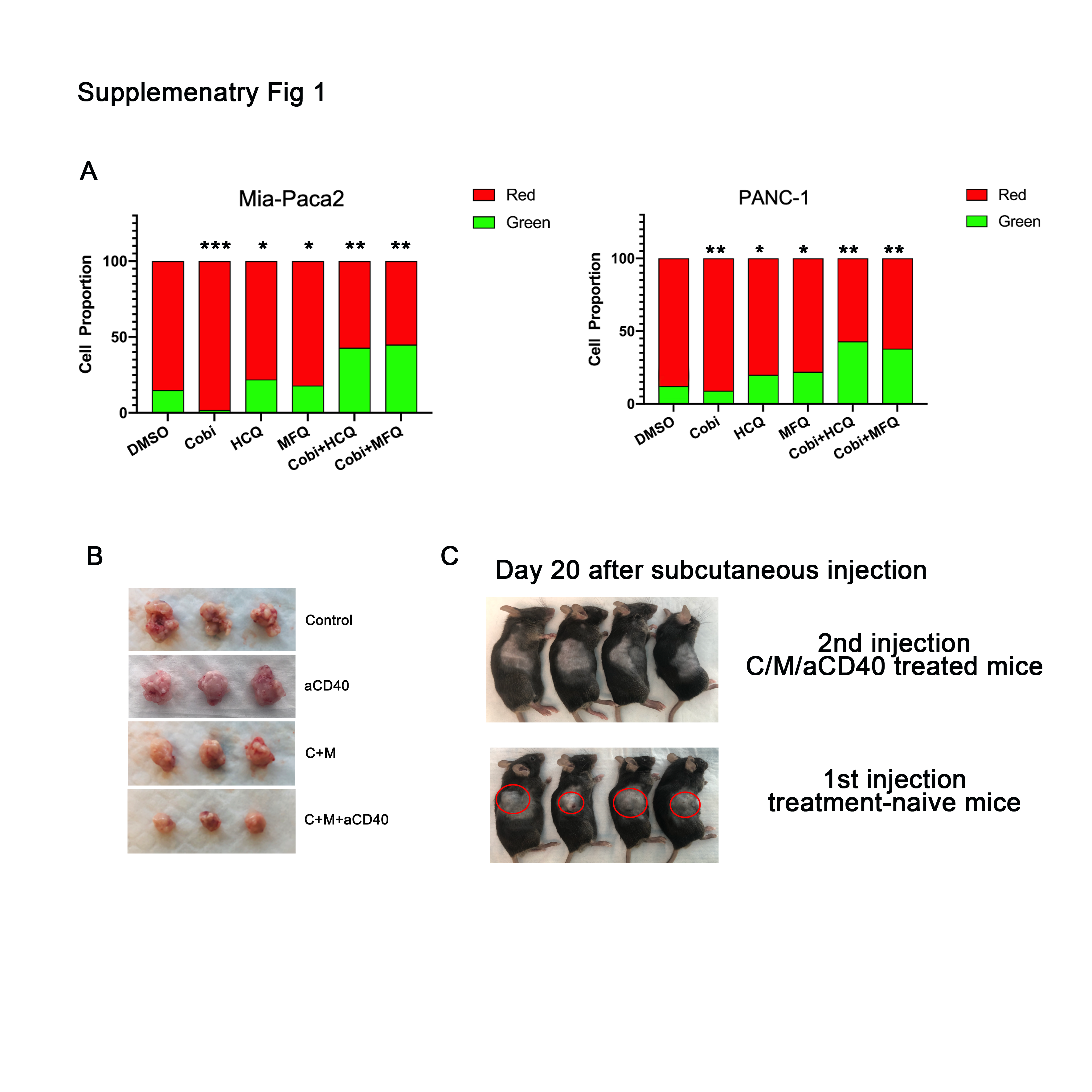

### Supplemental Fig2

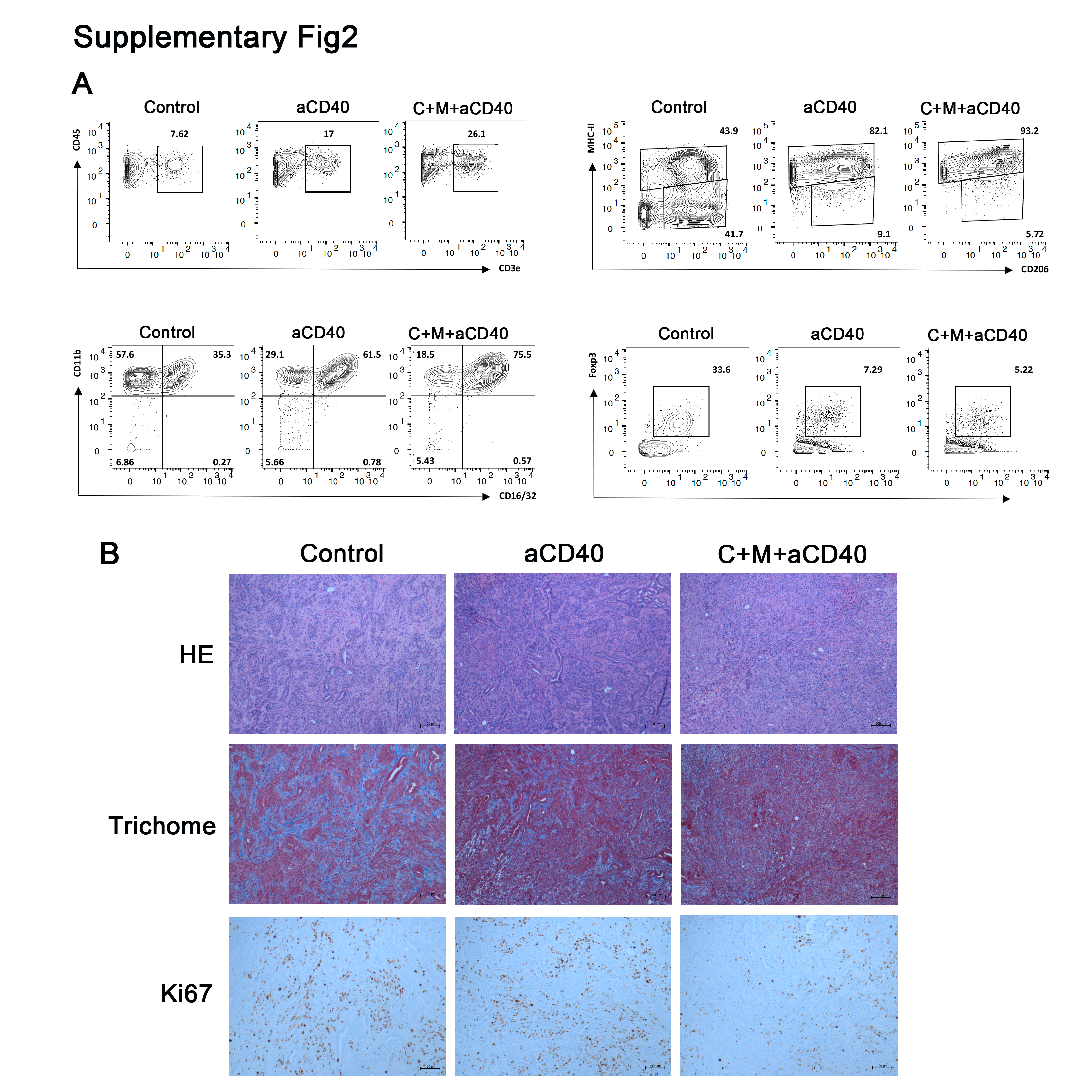

### Supplemental Fig3

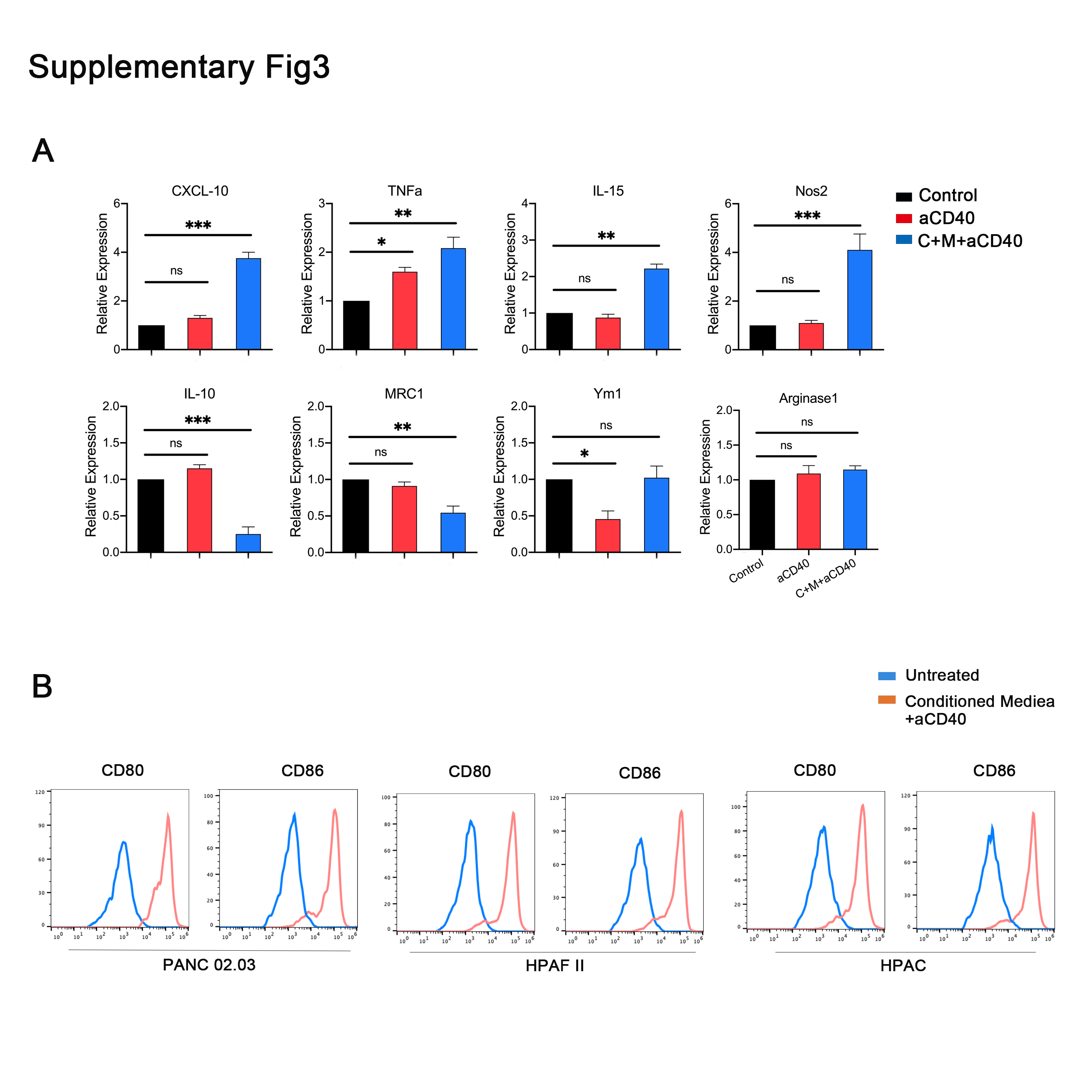

### Supplemental Fig4

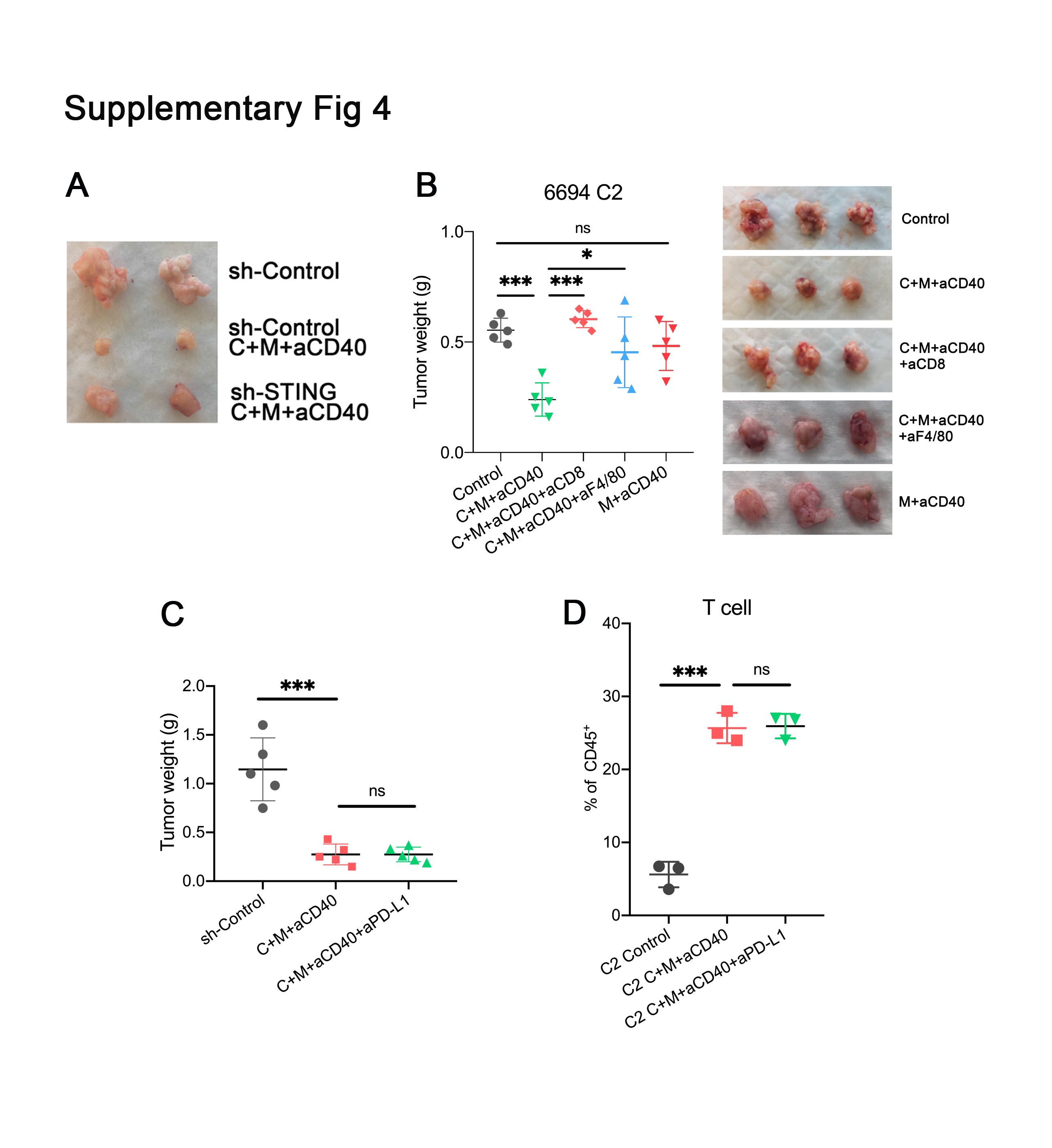

### Supplemental Fig5

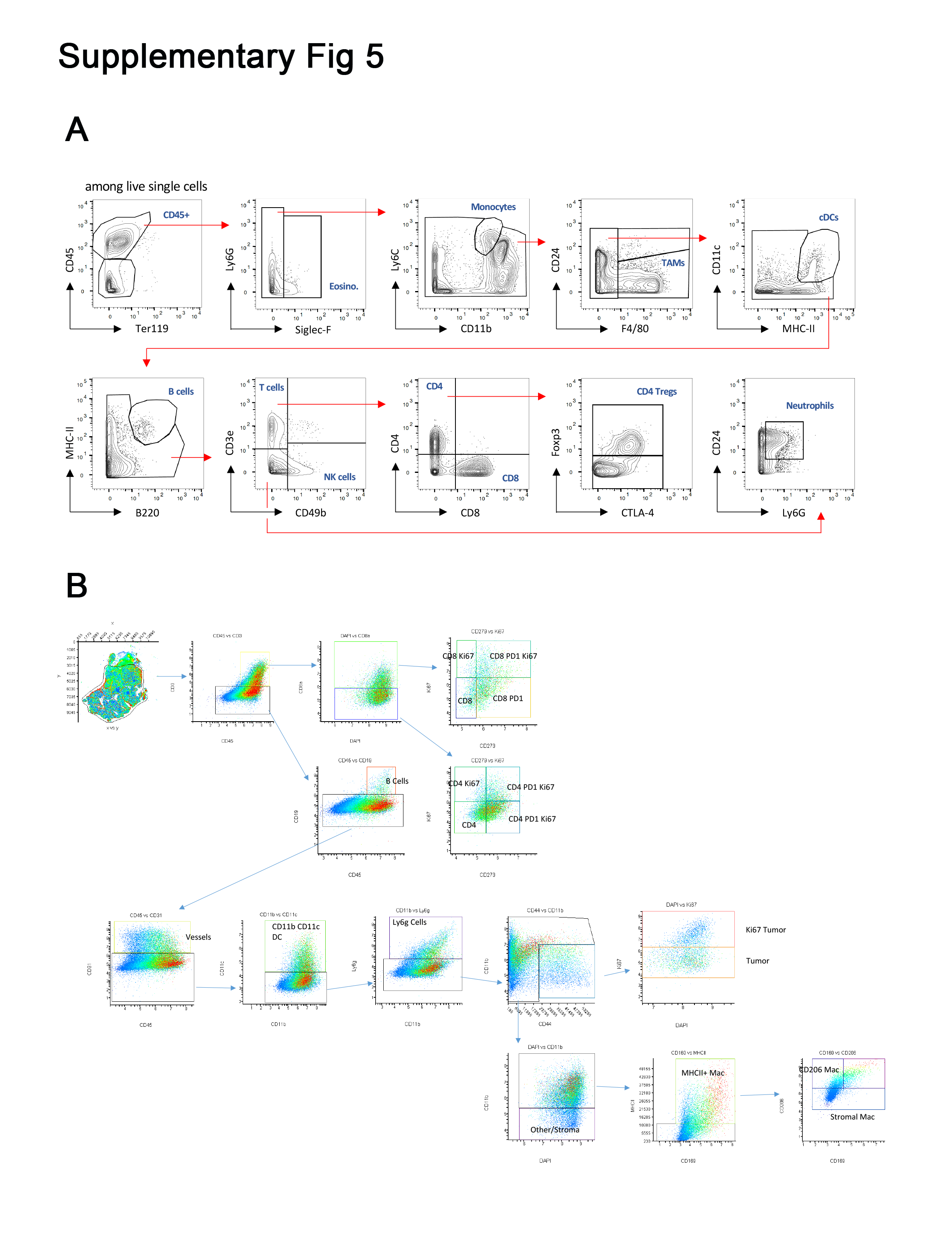
